# Modeling Zika Virus Spread in Colombia Using Google Search Queries and Logistic Power Models

**DOI:** 10.1101/365155

**Authors:** Mekenna Brown, Christopher Cain, James Whitfield, Edwin Ding, Sara Y Del Valle, Carrie A Manore

## Abstract

Public health agencies generally have a small window to respond to burgeoning disease outbreaks in order to mitigate the potential impact. There has been significant interest in developing forecasting models that can predict how and where a disease will spread. However, since clinical surveillance systems typically publish data with a lag of two or more weeks, there is a need for complimentary data streams that can close this gap. We examined the usefulness of Google Trends search data for analyzing the 2016 Zika epidemic in Colombia and evaluating their ability to predict its spread. We calculated the correlation and the time delay between the reported case data and the Google Trends data using variations of the logistic growth model, and showed that the data sets were systematically offset from each other, implying a lead time in the Google Trends data. Our study showed how Internet data can potentially complement clinical surveillance data and may be used as an effective early detection tool for disease outbreaks.

## 1 Introduction

The Zika Virus (ZIKV) disease is caused by a virus transmitted primarily by *Aedes* mosquitoes. The symptoms of ZIKV are typically mild and may include fever, rash, joint pain, and conjunctivitis (red eyes). These symptoms hardly warrant a visit to the hospital, and some infected persons may not exhibit any symptoms at all. The declaration of ZIKV to be a public health emergency was largely due to the correlation between ZIKV outbreaks and increased clusters of the neurological birth defect, Microcephaly [1]. In addition, ZIKV also poses a neurological threat to adults due to its link to the Guillain-Barre Syndrome [2]. Similar to other mosquito borne diseases, ZIKV appears to have a seasonal pattern. As of January 2018, the end of the mosquito season in many countries has slowed the spread, thus, the World Health Organization (WHO) has determined that ZIKV is no longer in a state of emergency. Nevertheless, ZIKV still poses a viable threat for upcoming seasons. Therefore, the WHO is developing a long-term response plan to minimize and ultimately prevent future outbreaks [3].

Traditional methods of case-counting during outbreaks result in long processing times, so case counts obtained using these methods typically lag behind real-time incidence by up to several weeks [4]. Recently, researchers have examined the potential use of Internet data streams to complement clinical surveillance data. In particular, various types of data have been used including Twitter, Google Trends, Facebook, Wikipedia, HealthMap, and others [5]. Internet data is appealing because it is updated frequently and is therefore expected to offer a near real-time data source for researchers and others to access.

Due to the prevalence of Internet usage, it is expected that physical phenomena would be expressed in Internet search patterns. If Internet searches of a disease are testament to an individual’s interest in that disease, it may be possible to quickly detect outbreaks and provide real-time information to health workers weeks before traditional methods do. Some attempts have been made to use real-time Internet search data to track outbreaks more effectively [5–7]. The goal of our study is to analyze the 2016 ZIKV outbreak in Colombia retroactively and determine whether Google Trends search queries might have served as a faster, up-to-date indicator of the ZIKV outbreak than traditional methods of data acquisition.

This paper is outlined as follows: Section 2 discusses previous approaches to modeling and predicting ZIKV and other diseases using Internet data and other related indicators. Sections 3 and 4 describe the types of data we analyzed in this study and our modeling approaches. The procedure of data correlation and the corresponding results are presented in sections 5 and 6. These are followed by discussions about the significance of our study and future research directions in sections 7 and 8.

## 2 Previous Approaches to Virus/Internet Modeling

A great deal of effort has been dedicated to finding efficient ways to track and predict the spread of infectious diseases over the years. The growing body of literature points to the use of Internet data as a potentially effective complimentary data set to inform models and subsequently predict disease spread. Successful modeling allows for early detection of outbreaks for up to one or two weeks, and that provides precious time for deploying educational programs aimed at mitigating disease spread as well as potentially managing resources in health care systems [8]. Numerous previous attempts for modeling and forecasting ZIKV and other diseases using Internet data have met some of success. Three examples are discussed in this section.

### 2.1 Google Flu Trends

Google Flu Trends (GFT), now defunct, was developed to make use of Internet search data to “now-cast” viral outbreaks of influenza [6]. This system monitored the volume of Google searches for key terms related to the virus and attempted to nowcast the number of cases of the disease [9]. The system functioned well during retroactive modeling, but since Internet behavior may change due to media coverage, it greatly over-predicted the number of cases for some seasons. Nevertheless, the study demonstrated the ability to leverage Internet search data to track influenza prevalence and provided valuable insights which led to the development of probabilistic and statistical models that describe the correlation between Internet data streams and disease outbreaks [10]. GFT was one of the first studies that showed promise in using Internet data to track diseases such as influenza.

### 2.2 Global Google Trends Study of ZIKV

A study closely related to ours is the work done by Teng *et al*. that retroactively compared cases of ZIKV calculated by WHO to publicly available Google Trends data for the search term “Zika” [7]. This study sought to determine if the global interest in Google searches for “Zika” correlated with the WHO’s ZIKV reported estimates. Since the study considered the global cumulative estimates of ZIKV cases and the world-wide Google searches for the term “Zika” it is considered a global study in scope. The study showed that the Internet searches for “Zika” and the total number of cases correlated well and that the Internet search data could be used to accurately predict the volume of ZIKV cases on a global scale. This method in comparing phenomenological models informs our attempt to use Google Search data for ZIKV on a departmental level in Colombia.

### 2.3 Antioquia Study

One study of ZIKV in Antioquia, Colombia by Chowell *et al*. focused on forecasting the growth of the disease using different logistic growth models [11]. The goal was to determine which model could forecast the transmissibility and final burden of the virus with the highest accuracy and predictive power. The model that worked the best is known as the Generalized Richards Model (GRM), and was able to forecast these factors accurately using data only from the first 30 weeks of the outbreak. On the contrary, the traditional logistic model was unable to reach this resolution even by the end of the outbreak. The work is useful for understanding the growth parameters of ZIKV, and we applied the GRM in our study to model the outbreak in all departments of Colombia.

The GFT and Global Google Trends studies of Zika were phenomenological models, correlating clinical surveillance to Internet search behavior. This phenomenological approach closely relates to the approach we took to produce the time series of ZIKV based on search data in Colombia. The previously published study of Antioquia was mechanistic in scope, with an attempt to understand the growth parameters of the virus. Our study applies the phenomenological approaches to correlate search patterns and ZIKV incidences quantitatively and utilizes the GRM to better compare the different data sets.

## 3 Sources of Data

Each of the aforementioned methods offers insight regarding successful uses of case incidence and Internet data. The goal of our study is to utilize these different data streams to accurately model existing data regarding the spread of ZIKV and determine if the Internet data has some level of predictive potential for clinical surveillance. The subject of our study is the 2016 ZIKV outbreak in Colombia for which complete and organized case data, published by the Colombian government, was obtained from a website accessed through Github [12]. We also used Internet search data that is publicly available on Google Trends [13].

### 3.1 Case Data

The case data we used is similar to the aforementioned Antioquia study [11]. The primary difference is that our data set covers all of the departments in Colombia (note that in Colombia “departments” refer to states). Case counts were originally divided into suspected cases, confirmed cases, and total cases [12]. For simplicity we considered only the total case counts for different provinces. All data is cumulative, collected over the first 37 weeks of 2016 during the height of the ZIKV outbreak in Colombia. The portion of data that was collected after this period was disregarded, as it had insignificant contributions to the total counts. One of our data sets, representing the cumulative case data in Tolima, Colombia, is plotted in Fig. 1. Note that some of the cumulative case counts contained obvious errors (e.g., cumulative totals dropped from one week to the next), which could have been caused by misreporting or typographical errors. For all municipalities, decreases in cumulative totals from one week to the next were set to 0. Precisely, for any week *t* and cumulative case count *C*(*t*), whenever *C*(*t* + 1) < *C*(*t*) or *C*(*t* +1) was missing, we set *C*(*t* + 1) = *C*(*t*). All non-decreasing cumulative totals were assumed to be correct. The slope of the trend approached zero toward the end of the ZIKV outbreak, indicating a steady decline in new cases. The growth in the cumulative counts bears a close resemblance to a general logistic growth pattern.

**Fig 1.**
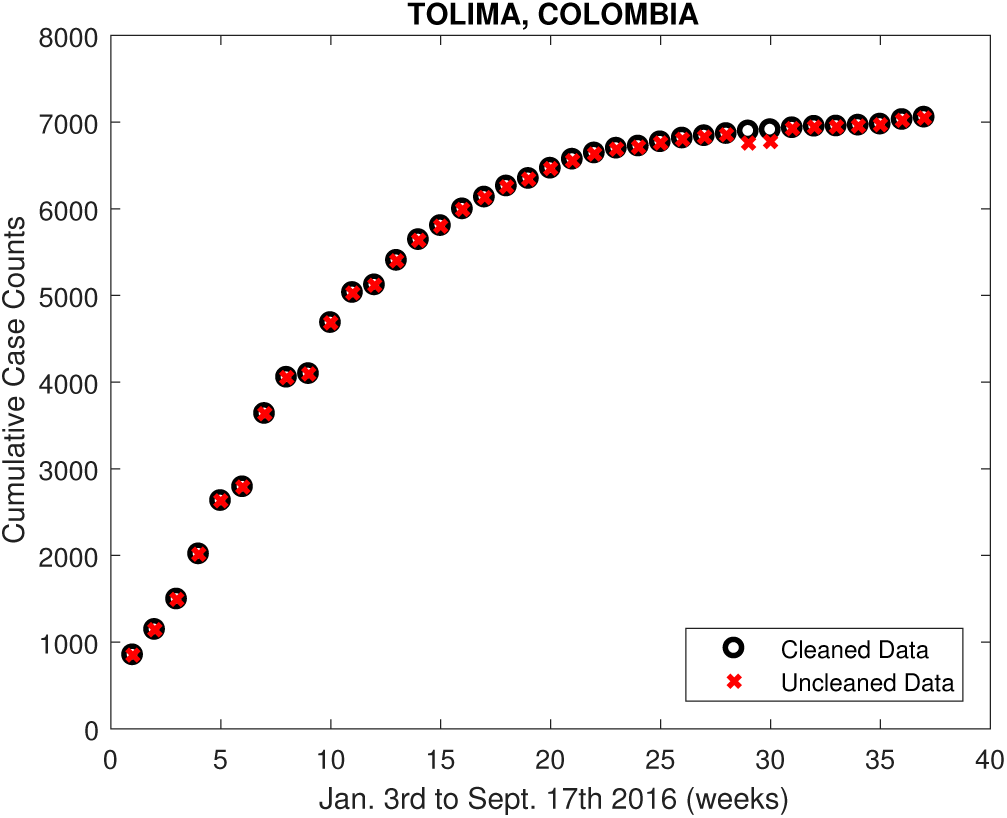
Cumulative case data versus time (in weeks) in Tolima, Colombia. The data was published publicly by the Colombian government in cumulative form on Github. The figure shows some of the cataloging errors (i.e., decreasing cumulative counts) we encountered when reviewing the data. Note that obvious errors wee removed at the municipality level of resolution and the cleaned data was subsequently used.

### 3.2 Google Trends Data

Our data from Google Trends [13] was taken in the period from March 1^st^, 2015 to February 2^st^, 2017, and was given in the form of cumulative relative frequency counts. Note that this time interval is larger than that of the case data and covers the entire ZIKV outbreak in the Google Trends data. Google Trends publishes the popularity of search terms and phrases in a particular department in weekly intervals. The search with the highest popularity over a selected time interval is set to 100, and all the other less popular searches are assigned values that are normalized with respect this value. These frequency counts are referred to in the literature as Google Trends Volumes (GTVs) [7]. We expect GTVs for certain search terms to correlate with the growth of ZIKV cases, so the two data streams should obey similar population growth models. We collected GTVs for the following ZIKV-related search terms in Spanish (English translation is included in parentheses): Zika, Zika Síntomas (Zika Symptoms), Virus Del Zika (Zika Virus), Sintomas Del Zika (Symptoms of Zika), and Zika Tratamiento (Zika Treatment). Data was collected at the departmental level in Colombia. To smooth out the noise in the data and to make direct comparisons to the cumulated case data, we also took the cumulative search volumes over the time interval of interest. The cumulative search volume counts appeared to match very well with what would be expected for a logistic growth model, just like the case data shown in Fig. 1. A representative example of our GTVs is shown in Fig. 2.

**Fig 2.**
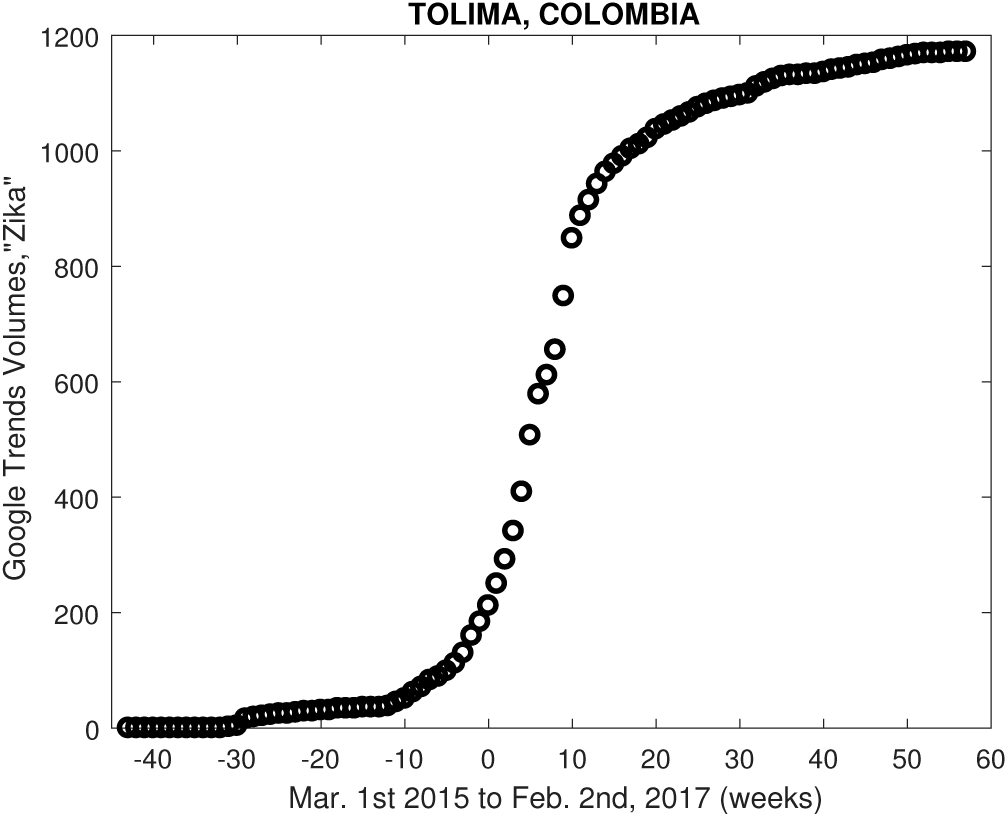
Cumulative GTVs in Tolima, Colombia versus time (in weeks). The data was originally published by Google Trends in weekly form, with the highest search frequency over the given time interval set to 100. To reduce the effect of weekly fluctuations, we cumulated the data, which decreased noise significantly.

## 4 Mathematical Modeling of Case and Google Trends Data

Based on the trends displayed by ZIKV case data and GTVs as shown in Figs. 1 and 2, we determined that a logistic type growth model should be adequate in terms of describing the growth rate of the cumulative ZIKV cases during the outbreak in Colombia. Two such models were considered in our study that targeted different data sets.

### 4.1 The Generalized Richards Model

The generalized Richards model (GRM) was used to find a best fit curve to the data using the least squares method. The GRM is a modification of the traditional logistic equation:

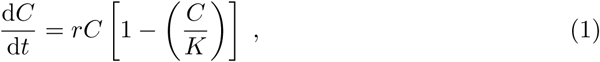

and was developed by Chowell *et al*. to more accurately predict Zika spread in Antioquia [11]. It has two extra parameters *p* and *a* that control the growth and decline rates of the model. The GRM is given by the differential equation

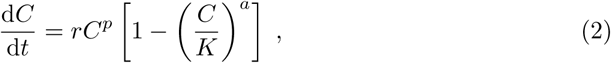

where *C*(*t*) represents the total case count as a function of time and *r* and *K* are logistic growth parameters. The special case of *a* = *p* =1 corresponds to Eq. (1). To simplify the computations needed for finding the best fit curve, we reduced the number of parameters in the GRM by a change of variables. Normalizing *C* with respect to the maximum case count *K*, i.e., the carrying capacity, the GRM can be expressed in the dimensionless form

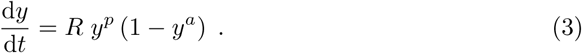

Here, *y*(*t*) = *C/K* is the normalized case count and *R* is related to the GRM parameters through *R* = *rK*^*p*−1^. The parameters *p, a*, and *R* are referred to in the literature as the deceleration growth, density dependence, and reduced unbounded growth rate, respectively [11].

Starting with a range of initial estimates of *p, a*, and *R*, we used the Levenberg-Marquardt Algorithm (LMA) in MATLAB to compute the optimal parameters that would produce the numerical solution of Eq. (3) that fit the data in the least squares sense. A Monte Carlo random sampling algorithm was then used to establish the confidence intervals of these optimal parameters. We employed a variant of the approach proposed by [14] using the Jacknife method instead of the Bootstrap method. In particular, 600 random samples of size 30 out of our 37 weeks of ZIKV case data were generated. These smaller subsets of the original data were each fitted using the LMA, yielding 600 sets of parameter estimates for *p, a*, and *R*. The average parameter values as well as their standard deviations were then calculated in order to establish the confidence intervals. Fig. 3 shows the best fit curves obtained using the full data set and the Monte Carlo process. The curves agreed extremely well with each other and also captured the patterns in the cumulative case data.

**Fig 3.**
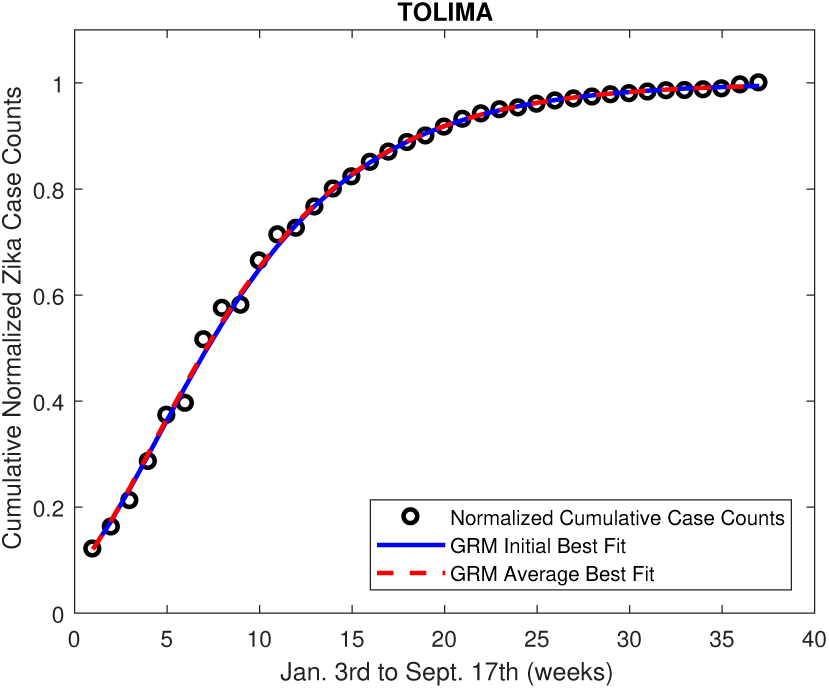
Best fit curves of the GRM to the Tolima, Colombia case data using the full case data set (red solid curve) and the Monte Carlo method (blue dotted curve). The circles represent the normalized case data. The initial fit was done once using all the data points. The average fit was determined by finding the best-fit curve from 600 random samples of size 30 using the Monte Carlo method.

A sampling of the parameter values *R* and *a* obtained using the Monte Carlo algorithm is given in Fig 4. These results imply that not only are these parameters highly sensitive to the initial guesses used in the LMA fitting procedure, but they are also impossible to be determined uniquely for a given data set. Sets of *R, a*, and *p* that differ significantly but give the same best-fit curve are given in Table 1, and the fits are plotted in Fig 5. This problem is not uncommon in the context of highly parameterized models, and is referred to in the literature as “parameter degeneracy” [15,16]. This occurs when two or more parameters in a model are affected by the same physical process in such a way that they cannot uniquely determined. It is common, however, for some combination of degenerate parameters to be uniquely determined [15]. In our case, the product *Ra* and 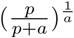 (the inflection point of the GRM) were always equal to 0.16 and 0.3295, respectively. Hence, even though the GRM parameters were degenerate in this case, certain combinations could be uniquely determined. Whether these results offer predictive value is beyond the scope of this study.

**Table 1.** A sample of average parameter values obtained using the Monte Carlo method. The data used for these fits was case data from Tolima, Colombia. *R* is the reduced unbounded growth rate, *a* is the density dependence, and *p* is the deceleration growth parameter. Note that *R* depends on the absolute unbounded growth rate (*r*), the deceleration growth parameter (*p*), and the total case capacity (*K*), as described earlier. Error measurements reflect a 95% confidence interval.

|  | Curve 1 | Curve 2 | Curve 3 |
| --- | --- | --- | --- |
| $R$ | $6.328 \pm 0.939$ | $10.597 \pm 1.187$ | $20.933 \pm 2.069$ |
| $a$ | $0.0253 \pm 0.0029$ | $0.015 \pm 0.0015$ | $0.0076 \pm 0.0006$ |
| $p$ | $0.888 \pm 0.00723$ | $0.893 \pm 0.0015$ | $0.896 \pm 0.0098$ |
| $Ra$ | $0.160 \pm 0.030$ | $0.159 \pm 0.024$ | $0.159 \pm 0.020$ |
| $(\frac{p}{p+a})^{\frac{1}{a}}$ | $0.329 \pm 0.0046$ | $0.329 \pm 0.0010$ | $0.329 \pm 0.0062$ |

**Fig 4.**
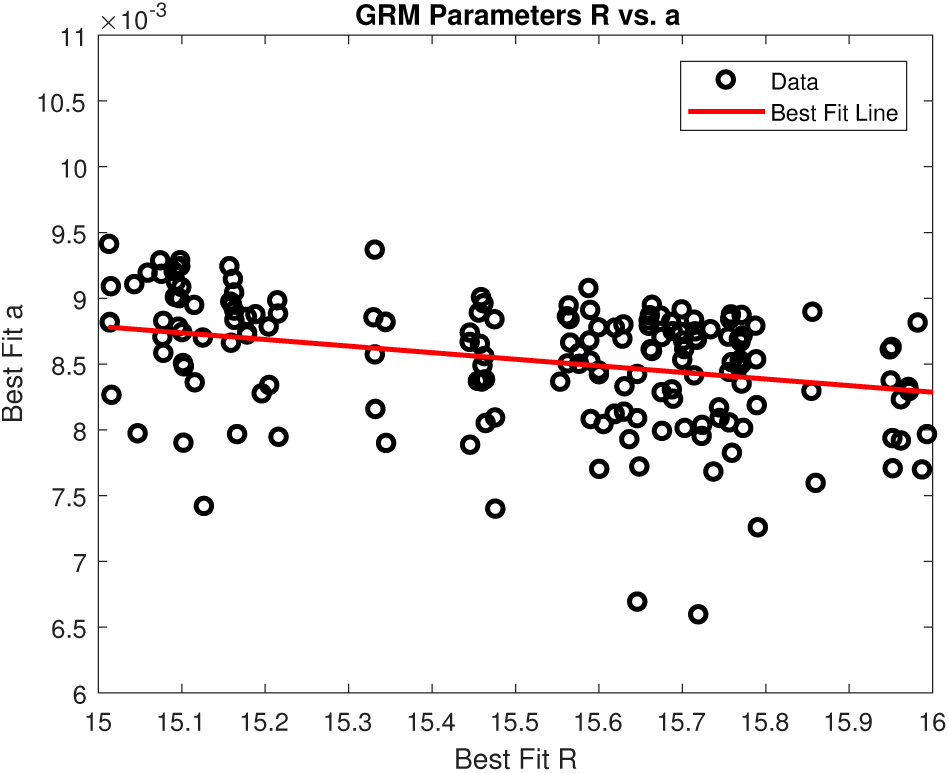
The computed values of *R* and *a* for all 600 random samples found using the Monte Carlo method on the data in Fig. 3. The best fit line that relates *R* and *a* is plotted in red. However, the global relationship is likely 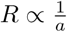, since we found *Ra* to be constant. This observation implies that *R* and *a* are free to vary according to some relationship while still producing an excellent best-fit curve.

**Fig 5.**
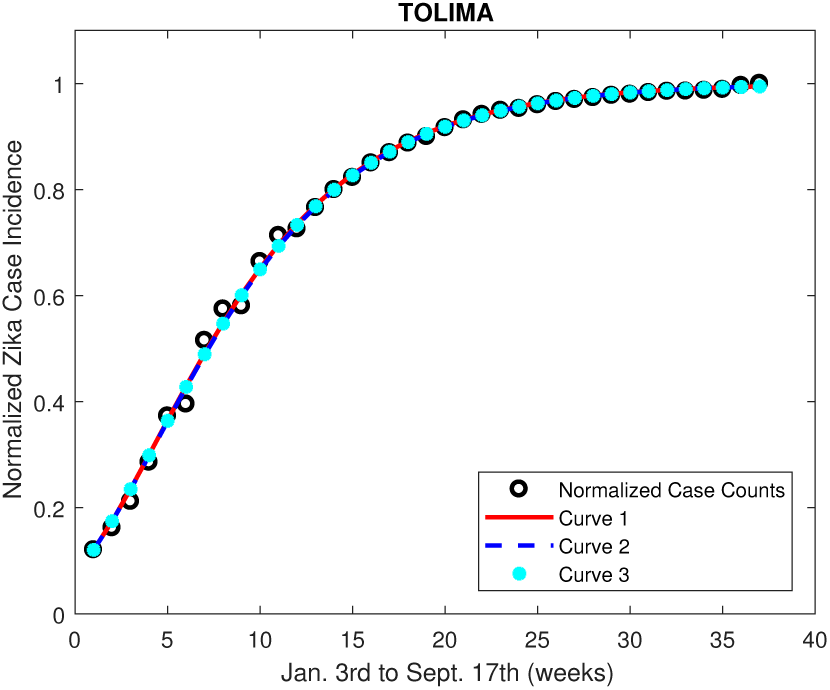
Normalized Tolima, Colombia data along with several best-fit curves obtained with different parameters. The actual parameters are given in Table 1. These parameters produce almost exactly the same best-fit curve to the data, suggesting that the parameters in the GRM cannot be uniquely determined using a best-fit analysis.

### 4.2 The Logistic Power Model

While the rate of change of the cumulated case counts approached zero toward the end of the outbreak, that was not the case in general for the GTVs. This is likely due to the presence of general search interest that was not a result of the outbreak itself. Since the GRM was not able to capture this general search interest, we used a different type of logistic growth model that can reflect the GTVs more accurately. The traditional logistic growth model, given by Eq. (1) has the general solution given by

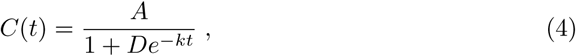

where *A* is the carrying capacity of the population, *k* is the unbounded growth rate, and 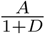 is the initial population. To account for non-uniform general search interest that does not obey a logistic growth pattern, we combined Eq. (4) with a power term to obtain

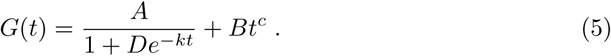

Here, *G*(*t*) denotes the cumulative GTVs. The added parameter c determines whether general interest is increasing (*c* > 1) or decreasing (0 < *c* < 1) and B is a parameter that weights the general interest component. We refer to this model as the Logistic Power Model (LPM). The interpretations of different model parameters are summarized in Table 2. The top panel of Fig. 6 shows the GTVs for the keyword “Zika” in Tolima, Colombia and the corresponding best fit curve in the form of the LPM and also how the latter compares to the GRM fit. The non-zero growth near the tail is well captured by the power term in the LPM. A general comparison between the traditional logistic model and the LPM is shown in the bottom panel where one can see the signature non-zero slope of the LMA near the tail. From the plots, it is clear that the LPM provides a more accurate description to the GTVs than the GRM does.

**Fig 6.**
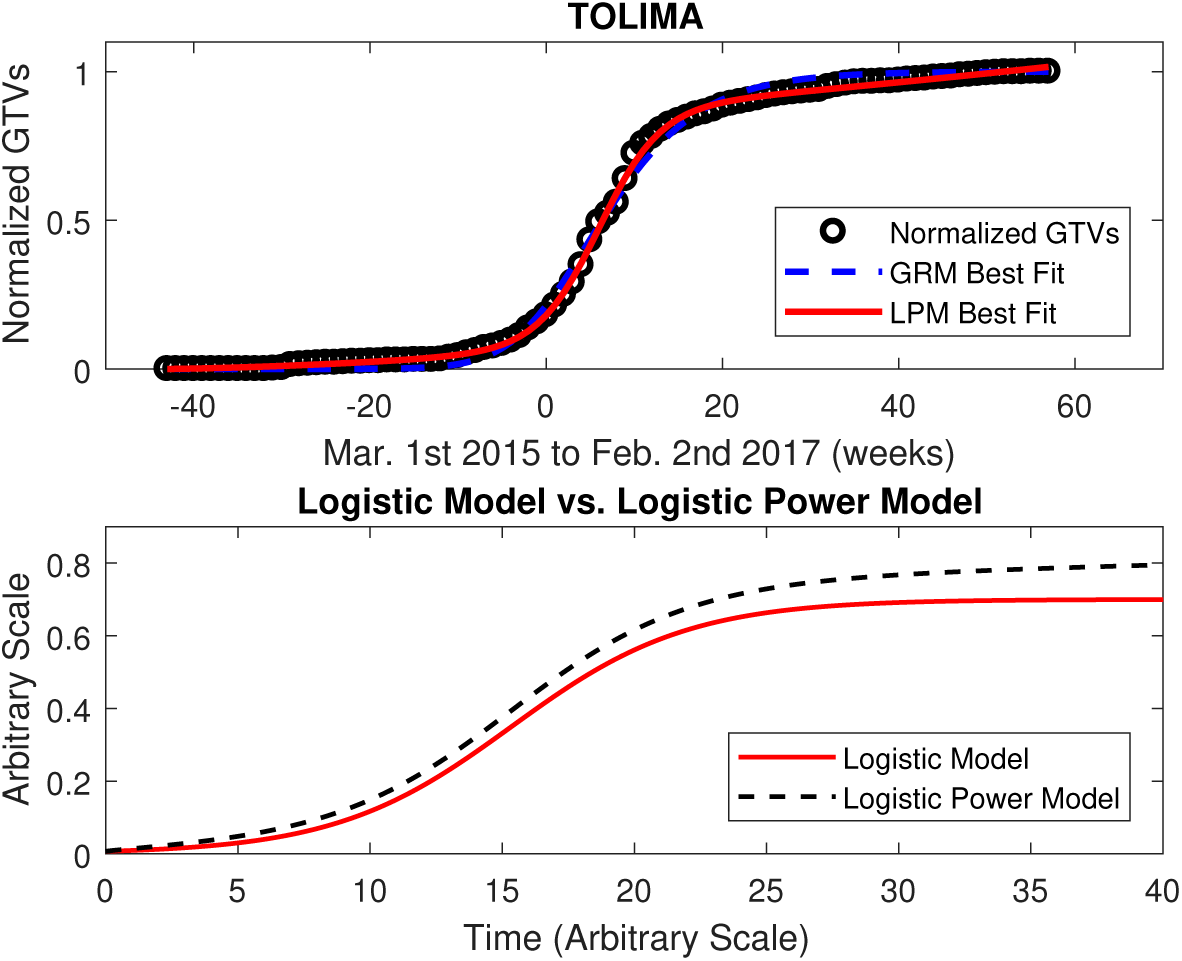
TOP: The best-fit curves to the GTVs for the keyword “Zika” in Tolima, Colombia using the logistic power model. For the LPM (black curve), the optimal parameters are *A* = 0.8330, *B* = 1.9101e — 04, *c* = 1.5264, *D* = 1.8050e + 06, and *k* = 0.2882. For the GRM (red curve), the fit parameters are *R* = 0.2177, *p* = 0.9597, and *a* = 0.9239. The results show that the LPM does a much better job of fitting the data than the GRM. One possible explanation is that the LPM accounts for general search interest, whereas the GRM does not. Note that in this case *c* > 1, which suggests an increasing rate of general search interest. BOTTOM: The form of the traditional logistic growth model as compared to that of the logistic power model. The parameters used for the logistic growth term of the LPM are the same as the parameters used for the traditional logistic growth model in this plot. The most interesting feature of the LPM is the non-zero slope at the tail end of the curve, which accounts for the fact that general search interest does not end at the conclusion of the outbreak.

**Table 2.** The parameters used in the normalized GRM Eq. (3) and LPM Eq. (5) are described here. All of these parameters, or variations thereof, have been used in previous modeling attempts except *B* and *c*, which are introduced to account for background search interest in the Internet data. Note that *R* depends on *p, K*, and *r* as described above.

| Parameter | Model used in | Description |
| --- | --- | --- |
| $a$ | Normalized GRM | Density dependence |
| $p$ | Normalized GRM | Deceleration growth |
| $R$ | Normalized GRM | Unbounded growth |
| $A$ | LPM | Carrying Capacity |
| $D$ | LPM | Normalization |
| $k$ | LPM | Unbounded growth rate |
| $B$ | LPM | Weight of general interest |
| $c$ | LPM | General Internet search interest |

To confirm statistically that the LPM estimates the Internet search queries more accurately than the GRM, we calculated the root-mean-square error (RMSE) for the best fit curves in each department in Colombia using GRM and LPM, respectively. A selection of RMSE values for the keyword “Zika” are given in Table 3. We consistently observed that the RMSE was higher for the GRM fits than for the LPM fits. We also performed a paired 2-sample *t*-test to compare the error in the two models, and obtained a test statistic of *t* = 8.805 and a *p*-value of 2.023 × 10^−9^. The statistical analysis performed here justifies choosing the LPM over the GRM to model the GTVs.

**Table 3.** The root-mean-square error (RMSE) for the GRM and LPM Google Trends fits for the search term “Zika” are given here. The error in the GRM fits was observed to be larger in nearly every case, which implies that the LPM performed better.

| Department | RMSE GRM | RMSE LPM | Difference |
| --- | --- | --- | --- |
| Antioquia | 0.0381 | 0.0271 | 0.0109 |
| Atlantico | 0.0193 | 0.0102 | 0.0090 |
| Bolivar | 0.0273 | 0.0163 | 0.0111 |
| Boyaca | 0.0386 | 0.0219 | 0.0166 |
| Caldas | 0.0439 | 0.0229 | 0.02010 |
| Caqueta | 0.0335 | 0.0159 | 0.0176 |
| Cauca | 0.0380 | 0.0246 | 0.0134 |
| Cesar | 0.0307 | 0.0131 | 0.0175 |
| Cordoba | 0.0289 | 0.0102 | 0.0186 |
| Cundinamarca | 0.0280 | 0.0164 | 0.0117 |
| Choco | 0.0362 | 0.0360 | 0.0002 |
| Huila | 0.0367 | 0.0099 | 0.0269 |
| La Guajira | 0.0366 | 0.0219 | 0.0147 |
| Magdalena | 0.0327 | 0.0168 | 0.0159 |
| Meta | 0.0298 | 0.0197 | 0.0100 |
| Narino | 0.0496 | 0.0206 | 0.0290 |
| Norte Santander | 0.0312 | 0.0110 | 0.0202 |
| Quindio | 0.0342 | 0.0284 | 0.0058 |
| Risaralda | 0.0394 | 0.0246 | 0.0149 |
| Santander | 0.0259 | 0.0208 | 0.0050 |
| Sucre | 0.0302 | 0.0122 | 0.0180 |
| Tolima | 0.0223 | 0.0091 | 0.0131 |
| Valle | 0.0200 | 0.0200 | 0.00 |
| Arauca | 0.0491 | 0.0225 | 0.0267 |
| Casanare | 0.0280 | 0.0141 | 0.0139 |
| Putumayo | 0.0242 | 0.0296 | -0.0054 |
| San Andres | 0.0371 | 0.0278 | 0.0093 |
| Amazonas | 0.0542 | 0.0455 | 0.0087 |

## 5 Correlating Data Streams

Previous work has demonstrated the existence of a correlation between Google Trends search volumes and ZIKV outbreaks [7]. Here, we show that ZIKV case data and related GTVs obey similar phenomenological models, so it is expected that they should be strongly correlated. More importantly, this correlation should reflect a common physical cause responsible for the behavior of both data sets (in this case, the ZIKV outbreak in Colombia). We hypothesize that ZIKV patients search for health-related information before seeking medical care. As such, clinical surveillance Internet data will be offset due to the significant lag between observation and reporting for clinical surveillance data when compared to the near-real time availability of Internet data streams.

To test this hypothesis, we probed the time dependence of the relationship between the GTVs and the case data using a time-series cross-correlation analysis. This approach is commonly used to identify time lags between independent and dependent physical variables [17,18]. The ZIKV case and GTVs were offset in time by an arbitrary time shift *τ* using the transformation

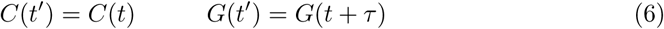

Here, *C*(*t*) represents the ZIKV case data for epidemiological (EPI) weeks 1 – 37 of 2016 [19] and does not change with *τ, G*(*t*) is the GTVs for the same time interval, and *t′* is the shifted time scale. The *t′* timescale remains the same for *C* but is shifted for *G* whenever *τ* ≠ 0, such that *G*(*t′*) and *C*(*t′*) are offset in real time. For example, for *τ* = 1 week, *C*(*t′*) and *G*(*t′*) correspond to the GTVs for EPI weeks 1 – 37 and 2 – 38, respectively. The value of *τ* was treated as a free parameter in our analysis. The Pearson’s r correlation between the offset data streams is then given by

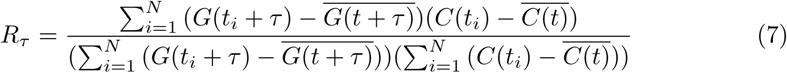

where *N* = 37 is the number of weeks of ZIKV data available and *R_τ_* is the correlation with time shift *τ* applied. Given the assumptions previously discussed, *C*(*t′*) and *G*(*t′*) should correlate better for some negative value of *τ* than at *τ* = 0. This should roughly correspond to the lead time of the Internet data. We refer henceforth to the value of *τ*, that maximizes *R_τ_* for some department and search term, as the Optimal Time Shift (OTS). Fig. 7 shows a comparison of case counts and GTVs for Tolima, Colombia (search term “Zika”), both normalized with respect to their values at EPI week 37, 2016. A small offset between the primary growth periods of the data sets is visible by inspection.

**Fig 7.**
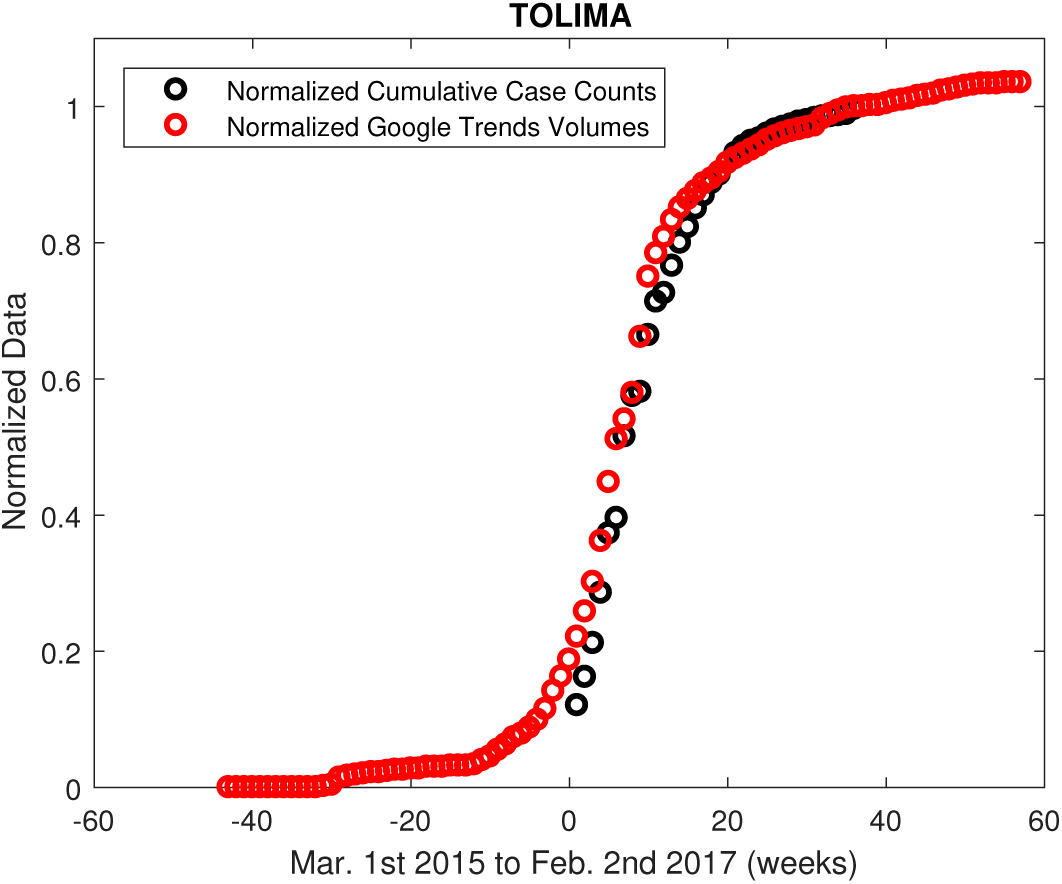
Normalized case counts and GTVs (search term “Zika”) for Tolima, Colombia, with no time shift. The data sets were both normalized with respect to their values at EPI week 37, 2016. A small offset in time between main growth periods of the data sets is visible when they are compared this way, suggesting a non-zero OTS.

*R_T_* was found for for –15 ≤ *τ* ≤ 15 (in weeks) in time steps of Δ*τ* = 1 week to determine OTS over a sufficiently large time-shift domain. This was done for each department and search term with available data. The graph of *R_τ_* vs. *τ* for Tolima, Colombia and search term “Zika” is given in Fig. 8.

**Fig 8.**
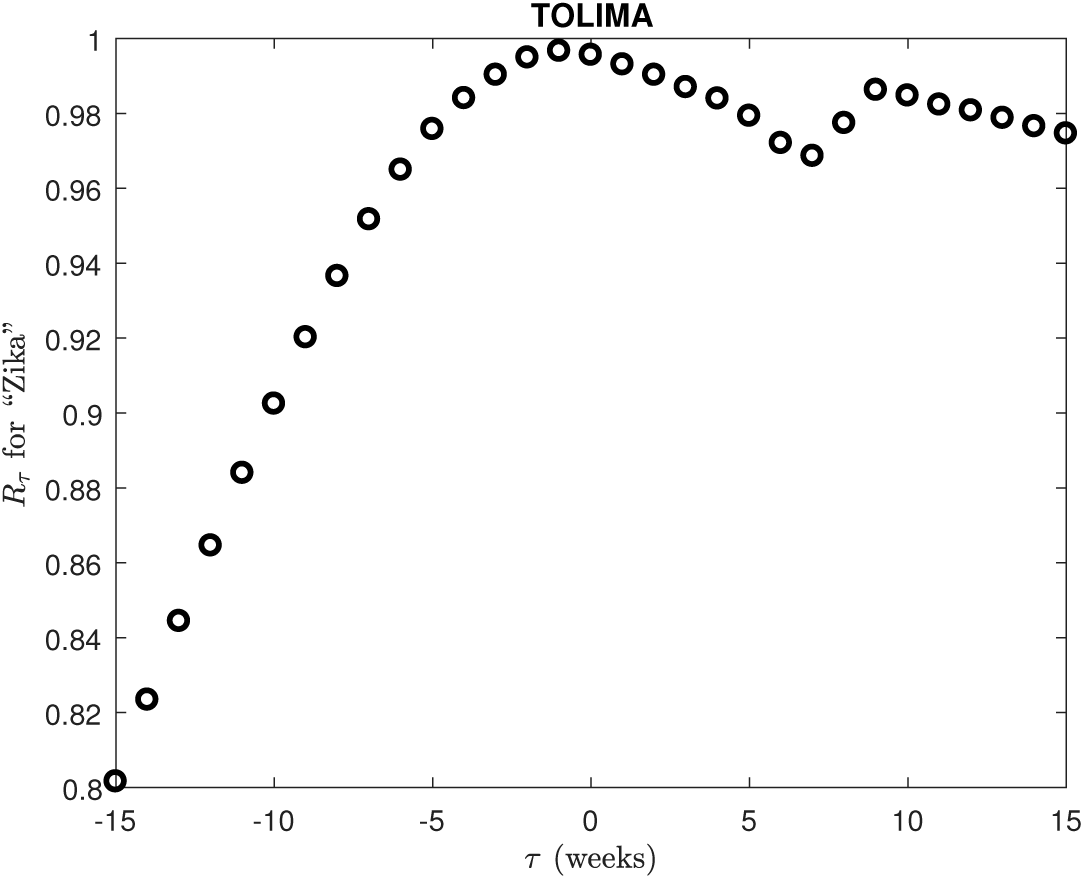
Correlation of 37 weeks of time-shifted case data and GTVs over the corresponding time interval for the search term “Zika”. The two data sets correlate extremely well near a time shift of 0, as expected. However, OTS is obtained at *τ* = – 1 week, suggesting a systematic time offset between the data sets. One possible interpretation of this offset is that Internet data is responding to the outbreak more rapidly than traditional case counting methods.

We expected to see some variability in OTS values across departments due to factors such as availability of hospitals, prevalence of Internet use, and other factors that affect both the cataloging of case data and frequency of searches. Furthermore, because ZIKV takes much more time to spread than news reports, departments geographically further from the location where the outbreak started may experience a larger gap between the initial rise in the frequency of Internet search and actual ZIKV cases. As a result, large variances in the OTS data should not be of concern, provided they are small enough to show that OTS < 0 on average. The average OTS 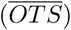, average optimal correlation 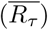 and standard deviation (*σ*(*OTS*)) across all departments with available data for 262 each search term is given in Table 4.

**Table 4.**
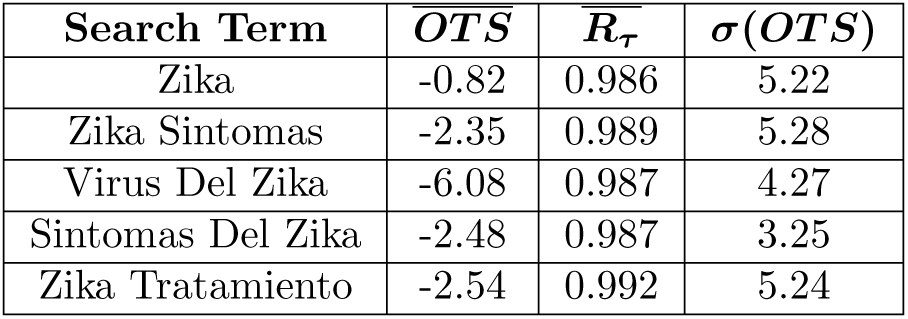
OTS averages. For all search terms considered, the OTS was negative, and the average optimal correlation 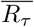 was very close to 1. However, *σ(OTS)* in each case was so large that a negative offset cannot be definitively identified. The focus of Section 6 is to try to resolve this issue. It is possible that contributions from departments with low cumulative case counts contributed anomalous results because they did not undergo true outbreaks during the time interval considered. This possibility prompted us to minimize the contribution from such departments in our analysis.

It is evident from Table 4 that negative time shifts tend to produce the best correlations. Still, the scatter in the data (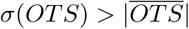 in all but one case) is large enough to suggest that the offset from 0 could be a random effect. One possible contributor to the observed scatter is the fact that many departments had relatively small ZIKV prevalence at the end of the outbreak. In these departments, the contribution to Internet search patterns by the outbreak may have been washed out by contributions from other sources. To account for this effect, we found 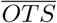 for each search term weighted by the percentage of the department populations that had been infected as of EPI week 37, 2016. These results are given in Table (5).

**Table 5.** OTS Averages with ZIKV Prevalence Weighting. OTS averages weighted by the total percentage of ZIKV cases as of EPI Week 37, 2016 in each department. Significant improvement was only seen for the search term “Zika”, which displayed a much more negative OTS and a smaller *σ*(*OTS*). The other search terms had either similar or less favorable results. For the phrase “Zika Síntomas”, the department of North Santander had an OTS value of +15, which is clearly a statistical outlier. With this data point removed, 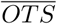 and *σ*(*OTS*) were –3.47 and 2.95, respectively. “Zika Tratamiento”, which provided the least conclusive results, had the fewest available data points (11).

| Search Term | $OTS$ | $\overline{R_\tau}$ | $\sigma(OTS)$ |
| --- | --- | --- | --- |
| Zika | -3.87 | 0.991 | 3.79 |
| Zika Sintomas | -1.38 | 0.991 | 7.14 |
| Virus Del Zika | -5.85 | 0.989 | 4.84 |
| Sintomas Del Zika | -2.77 | 0.989 | 3.41 |
| Zika Tratamiento | -1.31 | 0.993 | 5.24 |

Even with the weighted averages, *σ*(*OTS*) remains large in every case. This seems to be due to a combination of statistical outliers (see caption of Table 5) and the small size of the available data sets (“Zika Tratamiento” displayed the worst results, and had the smallest number of data points). Our analysis of the raw data suggests a systematic time offset between the data streams, but cannot show conclusively that one exists. In the next section, we make use of the phenomenological modeling results from Section 4 to try to resolve this situation.

## 6 Correlating Best Fit Curves

Even the best data sets available to us were subjected to sources of noise that may hamper an accurate investigation of intrinsic trends. For example, the normalization procedure used on the GTVs by Google Trends makes them somewhat vulnerable to quantization error at weekly intervals, which is itself cumulated when the data is aggregated. Both data types may be affected by cataloging errors and odd statistical fluctuations resulting from small sample sizes. These error sources, taken together, may have been responsible for some of the scatter observed in the correlation results in the previous section.

To circumvent these difficulties and correlate intrinsic data trends more accurately, we used the phenomenological modeling results from Section 4 to refine our correlation analysis. We did this by applying the same correlation procedure described in Section 5 to the best fit curves obtained in Section 4, with the GRM used to model the case data and the LPM used to model the GTVs. The best fit curves used in this section were computed over same time intervals from Section 3 in time-steps of *δt* = 0.1 weeks. As before, –15 ≤ *τ* ≤ 15 weeks, this time with Δ*τ* = 0.1 weeks. Because both models fit the data very well, correlating the fits should accurately relate the overall trends in the data with most of the noise filtered out. Henceforth, we refer to OTS values obtained by correlating the best fit curves obtained using the GRM and LPM as Model Optimal Time Shifts (MOTS).

The same correlation data shown in Fig. 8 for Tolima, Colombia is given again in Fig. 9, this time with the correlation of the best fit curves shown in the same plot. In this case, OTS occurs slightly earlier, and the behavior of *R_τ_* for *τ* > 0 is nearly monotonic. This behavior seems more reasonable because a local or global maximum in the positive time shift domain would imply a negative Internet lead time.

**Fig 9.**
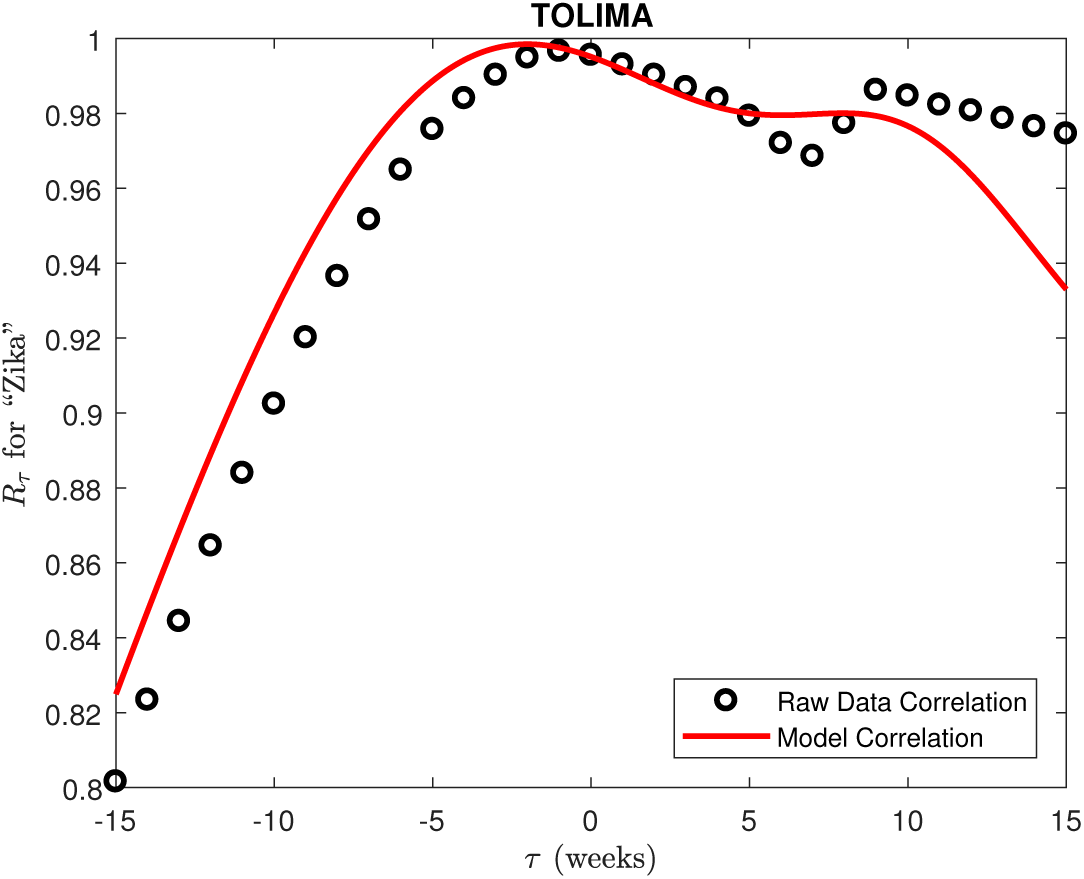
The same correlation data from Fig. 8, with best-fit curve correlation data added. The correlation peaks further to the left of 0, and the behavior to the right of 0 is nearly monotonic, unlike the raw data correlation.

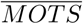 and *σ*(*MOTS*) for all search terms are given in Table (6). The systematic negative offset is much more prominent here than in Table 4. Moreover, 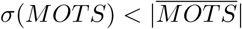 in most cases, making these results much more promising. To further verify this result, we repeated the weighting scheme used in Section 5 with the MOTS data. These results are given in Table 7.

**Table 6.** MOTS Averages. Optimal time shifts found using the best fit curves were more consistently negative than in Table 4. All the GRM and LPM fits used to calculate MOTS did a good job of capturing the data trends. These results provide more convincing evidence that the intrinsic outbreak-dependent features of the data are offset from each other in time.

| Search Term | $\overline{MOTS}$ | $\overline{R}_\tau$ | $\sigma(MOTS)$ |
| --- | --- | --- | --- |
| Zika | -4.42 | 0.993 | 3.11 |
| Zika Sintomas | -4.15 | 0.997 | 2.40 |
| Virus Del Zika | -6.42 | 0.995 | 4.82 |
| Sintomas Del Zika | -3.68 | 0.996 | 2.40 |
| Zika Tratamiento | -4.97 | 0.995 | 3.43 |

**Table 7.** MOTS Averages with ZIKV Prevalence Weighting. Results of curve correlation with weighted average. As in Section 5, the weighting scheme used did not make a significant difference to the results.

| Search Term | $\overline{MOTS}$ | $\overline{R_\tau}$ | $\sigma(\overline{MOTS})$ |
| --- | --- | --- | --- |
| Zika | -4.62 | 0.996 | 2.51 |
| Zika Sintomas | -4.63 | 0.998 | 2.60 |
| Virus Del Zika | -6.06 | 0.997 | 4.92 |
| Sintomas Del Zika | -3.38 | 0.997 | 2.54 |
| Zika Tratamiento | -4.33 | 0.995 | 3.00 |

Because the OTS and MOTS statistics depend partially on the behavior of the case data, they should be at least weakly correlated to physical variables affecting the spread of ZIKV. Temperature and other environmental factors are known to heavily influence the spread of ZIKV [20]. We investigated humidity and temperature because both vary widely across Colombia, and department-level data was readily available. For each physical variable, we measured two statistics for each department in Colombia: the average value from January to September, and the month of the year during which the peak value was measured. This data was compared to OTS and MOTS averaged across all available search terms for each department (denoted **OTS** and **MOTS** to differentiate between the associated averages across all departments for each search term) to determine if any relationship exists. The results are given in Figs. 10 and 11.

**Fig 10.**
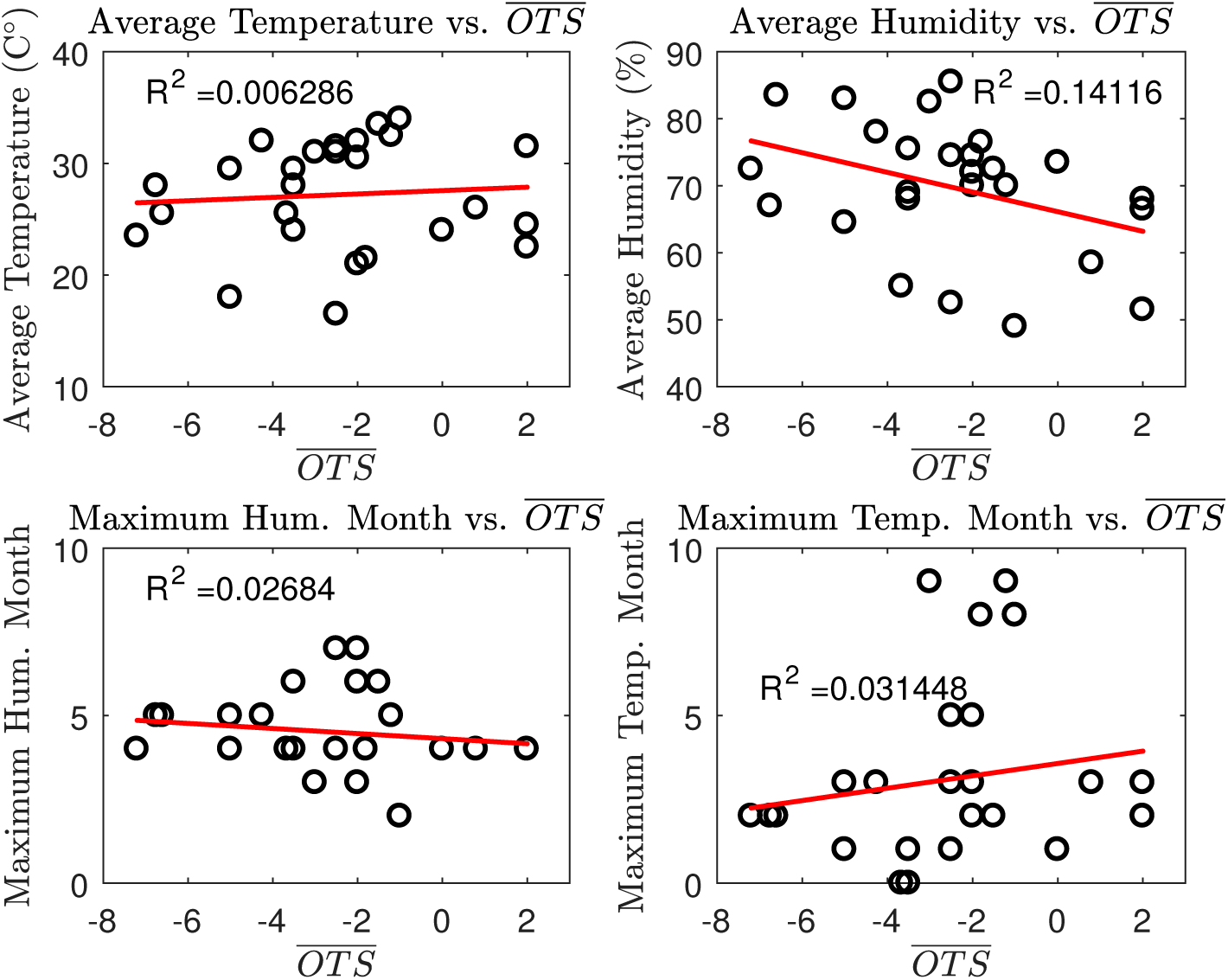
Best-fit lines (solid) for average temperature, average humidity, maximum humidity month, and maximum temperature month vs. **OTS** for departments with available weather data (between Jan. and Sept. 2016) and GTVs (data in black). The *R*^2^ values for each plot are 0.0063, 0.1414,0.027, and 0.031, respectively. A weak correlation to average humidity is suggested by the data, but otherwise no definitive relationships can be observed.

**Fig 11.**
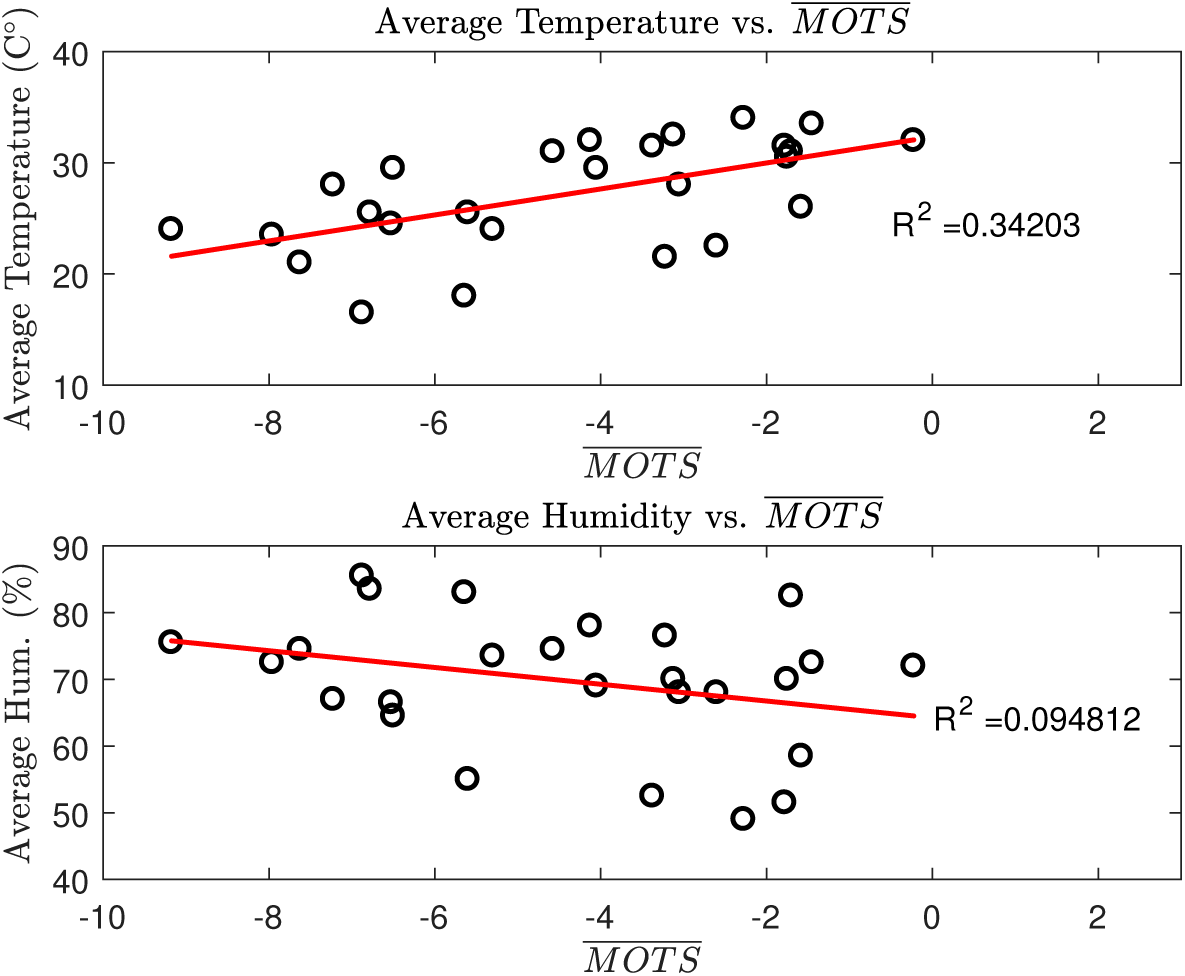
Best-fit lines (solid) for average temperature and average humidity (Jan. - Sept. 2016) vs. **MOTS** (data in black). The average humidity correlation observed in Table 10 is weaker here (*R*^2^ = 0.095), suggesting that there may be no intrinsic relationship. However, a notable correlation to average temperature is observed (*R*^2^ = 0.342), helping to explain the large scatter observed in the **MOTS** data.

Both **OTS** and **MOTS** appear to be weakly anti-correlated to average humidity, with the **OTS** correlation being slightly stronger. Humidity high month and temperature high month do not correlate to either **OTS** or **MOTS**. However, the relationship to average temperature is much more interesting. A small, if any, relationship between average temperature and **OTS** can be inferred from this data.

However, a notable positive correlation (*r* = 0.584, *R*^2^ = 0.342) exists between average temperature and **MOTS**. This makes sense because ZIKV is known to spread more effectively at higher temperatures [20]. Departments experiencing faster ZIKV spread would experience outbreaks that develop more rapidly, which may in turn reduce the lag time of reported case data with respect to Internet data. This result suggests a negative relationship between Internet lead time and average temperature. Table 8 displays temperature and humidity averages alongside **OTS** and **MOTS** for all departments with available data. The departments are organized by increasing average temperature.

**Table 8.** Temperature and Humidity Summary Table. A summary of **OTS, MOTS**, average temperature and average humidity by department. We also include MOTS for the search term “Zika”, which had generally the best data resolution. The positive correlation between **MOTS** and average temperature is clear, while **OTS** does not appear to correlate well to either variable.

| Department | OTS | MOTS | MOTS (“Zika”) | Avg. Temp. (C°) | Avg. Hum. (%) |
| --- | --- | --- | --- | --- | --- |
| Boyaca | -2.5 | -6.875 | -7.9 | 16.5 | 85.5 |
| Cudinimarca | -5 | -5.64 | -3.7 | 18.0 | 83 |
| Narino | -2 | -7.625 | -9.1 | 21.0 | 74.5 |
| Antioqua | -1.8 | -3.22 | -4.4 | 21.5 | 76.5 |
| Caldas | 2 | -2.6 | -5 | 22.5 | 68 |
| Santander | -7.2 | -7.96 | -7.6 | 23.5 | 72.5 |
| Putumayo | 0 | -5.3 | -5.1 | 24.0 | 73.5 |
| Cauca | -3.5 | -9.175 | -12 | 24.0 | 75.5 |
| Risaralda | 2 | -6.525 | -7.4 | 24.5 | 66.5 |
| Valle | -6.6 | -6.78 | -6.2 | 25.5 | 83.5 |
| Tolima | 0.8 | -1.58 | -2 | 26.0 | 58.5 |
| Quindio | -3.67 | -5.6 | -8.8 | 27.0 | 51.5 |
| Meta | -6.75 | -7.225 | -6.4 | 28.0 | 67 |
| Caqueta | -3.5 | -3.05 | -0.3 | 28.0 | 68 |
| Casanare | -5 | -6.5 | -7.9 | 29.5 | 64.5 |
| Arauca | -2 | -1.75 | -5.9 | 30.5 | 70 |
| Magdalena | -2.5 | -4.58 | -4.2 | 31.0 | 74.5 |
| Amazonas | -3 | -1.7 | -1.7 | 31.0 | 82.5 |
| Norte Santander | 2 | -1.78 | -2.7 | 31.5 | 51.5 |
| Huila | -2.5 | -3.38 | -2.4 | 31.5 | 52.5 |
| Guajira | -3.5 | -4.05 | -2.6 | 32.0 | 69 |
| Bolivar | -2 | -0.225 | -0.7 | 32.0 | 72 |
| Cordoba | -4.25 | -4.125 | -2.6 | 32.0 | 78 |
| Atlantico | -1.2 | -3.12 | -1.3 | 32.5 | 70 |
| Sucre | -1.5 | -1.45 | -2.5 | 33.5 | 72.5 |
| Cesar | -1 | -2.28 | -1.8 | 34.0 | 49 |

The observed relationship between MOTS and average temperature also helps explain why *σ*(*MOTS*) remained large even when the best-fit curves were correlated. The scatter in the data may well have been at least partially a product of temperature dependence across departments. Furthermore, the significant improvement in the temperature correlation suggests that scatter in the raw data may be hiding the intrinsic temperature relationship. OTS likely produced a poor correlation because poor resolution in the raw data produced erroneous correlation results for many departments. The modeling procedures used to produce MOTS appear to have at least partially resolved that issue. What is most important is that MOTS was consistently negative for all search terms, and 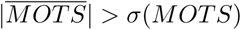 for all search terms when statistical outliers were neglected. Hence, a systematic offset between reported ZIKV case and GTV data streams is evident when phenomenological models are used in conjunction with a correlation analysis.

## 7 Discussion

As mentioned in section 4, ZIKV case and Internet search data are accurately described with variation on the logistic growth function. The model that effectively describes the case data is the GRM. The model that most effectively described the Internet data is the LPM, which adds a variable power law to the logistic growth function to account for general search interest. This model may prove useful in future studies of Internet trends. Note that the parameters within the GRM and LPM were not analyzed for significance, but were optimized for most accuracy in representing the data. Future work may focus on better understanding the physical implications of the parameters.

Although we were not able to uniquely determine the GRM parameters for the case data, the results were still interesting for various reasons. First, it was clear that the model fit the data well, regardless of the particular parameter values produced by the least squares method. This serves as strong evidence that cumulative case counts did obey a generalized population growth model, as demonstrated in previous work [11]. Moreover, we found that some physically relevant quantities, such as the point of maximum growth (i.e., inflection point), are invariant regardless of the individual parameter values. Thus, despite our difficulties, we were able to extract some interesting results from our efforts to model the case data.

Direct and indirect examination of the correlation between data streams provided evidence supporting a systematic time offset between reported ZIKV case and GTVs during the 2015 – 2016 Colombia outbreak. Preliminary investigation of the OTS statistic revealed that the data streams tend to correlate better when ZIKV case data is shifted to an earlier time interval. When the phenomenological models discussed in section 4 are incorporated, the time offset becomes much more pronounced. One possible interpretation of this trend is that Internet search patterns in Colombia reacted to the ZIKV outbreak more quickly than did the actual reported case data obtained using traditional methods. If this is the case, Internet search data repositories such as Google Trends may have the potential to provide health professionals earlier indication of outbreaks of ZIKV and as such, complement traditional methods.

Our analysis of OTS and MOTS suggests that the time delay between case and Internet data may depend heavily on a variety of geographical factors. As previously mentioned, it is possible that news reports about ZIKV triggered by the initial outbreak traveled throughout Colombia much faster than ZIKV, prompting Internet searching interst in departments that did not themselves experience outbreaks until weeks later. This effect may be at least partially responsible for the time offset. Our results suggest that the strong dependence of ZIKV spread on average temperature may have an effect on the offset as well. It will be left to future research to probe these dependencies more thoroughly.

## 8 Conclusions

Two conclusions can be drawn based on the results of our study. First, reported Zika case data and Zika-related GTVs in Colombia can be accurately modeled using logistic type growth models. The generalized Richard’s model (GRM) provides an accurate description of the cumulative case counts during the outbreak, while the Google Trends data can best be modeled using the logistic power model (LPM) that accounts for background searches unrelated to the Zika outbreak. Second, we concluded that Google Trends data is a potential early indicator of the spread of ZIKV compared to traditional case-counting methods. Our results yielded strong evidence for a systematic offset between the two data streams that suggests a lead time in Internet data. The calculated lead time varied widely by department, and preliminary investigation reveals that temperature may be a significant contributing factor to this observed variability. These results imply that GTVs has the potential to serve as an early detection tool for health agencies. Further research may investigate the dependence of the lead time more thoroughly.

## 9 Acknowledgements

The authors acknowledge support from PIC Math, a program of the Mathematical Association of America (MAA) and the Society of Industrial and Applied Mathematics (SIAM), and the National Science Foundation (NSF grant DMS-1345499). SYD and CAM were supported by Los Alamos National Laboratory (LANL). CAM was partially funded by the NSF SEES grant CHE-1314029 and LANL LDRD Director’s Postdoctoral Fellowship. LANL is operated by Los Alamos National Security, LLC for the Department of Energy under contract DE-AC52-06NA25396. Approved for public release: LA-UR-18-23845. The funders had no role in study design, data collection and analysis, decision to publish, or preparation of the manuscript.

